# A robust Reeb graph model of white matter fibers with application to Alzheimer’s disease progression^⋆^

**DOI:** 10.1101/2022.03.11.482601

**Authors:** S. Shailja, Scott T. Grafton, B. S. Manjunath

## Abstract

Tractography generates billions of complex curvilinear fibers (streamlines) in 3D that exhibit the geometry of white matter pathways. Analysis of raw streamlines on such a large scale is time-consuming and intractable. Further, it is well known that tractography computations produce noisy streamlines, and this in turn severely affect their use in structural brain connectivity analysis. Prompted by these challenges, we propose a novel method to model the bundling structures of streamlines using the construct of a Reeb graph. Three key parameters in our method capture the geometry and topology of the streamlines: (i) *ϵ* – distance between a pair of streamlines in a bundle that defines its sparsity; (ii) *α* – spatial length of the bundle that introduces persistence; and (iii) *δ* – the bundle thickness. Together, these parameters control the robustness and granularity of the model to provide a compact signature of the streamlines and their underlying anatomic fiber structure. We validate the robustness of the bundling structure using synthetic and ISMRM datasets. Next, we demonstrate the potential of this approach as a tool for efficient tractogram comparison by quantifying the fiber densities in the progression of Alzheimer’s disease. Our results on ADNI data localize the maximal bundles of various brain regions and show a significant depletion in the fiber density as Alzheimer’s disease progresses. The source code for the implementation is available on GitHub.

## 1 Introduction

Diffusion Tensor Imaging (DTI) is crucial for structural brain analysis to identify white matter alterations in many neurological applications such as aging, development, and disease [5,10,24]. Using tractograms for the study of structural connectomics and analyses of neurological diseases is challenging due to the complex structure of the white matter anatomy and the large number of streamlines generated by DTI tractography. Hence, quantitative analyses of streamlines is of great interest [31]. The current research on DTI streamlines analyses can be classified into two broad categories — connectome and tract-specific. While connectome studies properties of connectivity of brain regions, the tract-based analysis is focused on clustering anatomically known sets of fibers for a particular application. The common thread in both of these methods is that they use dimensionality reduction to represent the billions of streamlines to yield feasible computational models. Connectome analysis [8,30] usually generates a connectivity matrix by partitioning the brain cortex into a limited set of regions that are derived from anatomical or computational brain atlases. The streamlines within each brain region are lumped into a single node in this representation to model their interconnections as shown in Fig. 1. Hence, the dimension of a connectivity matrix is equal to the number of regions of interest (usually ∼ 100). Connectivity matrices provide measures of structural connectivity of the entire brain [11,19]. On the other hand, tract-specific analysis methods [12,29] cluster the massive number of tractography fibers to study domain-specific problems. Various fiber clustering methods (spectral clustering, hierarchical clustering, k-nearest neighbors) and similarity metrics (covariance matrix, Hausdorff [6], Mahalanobis distance [13]) have been studied to identify homogeneous clusters [9].

**Fig. 1.**
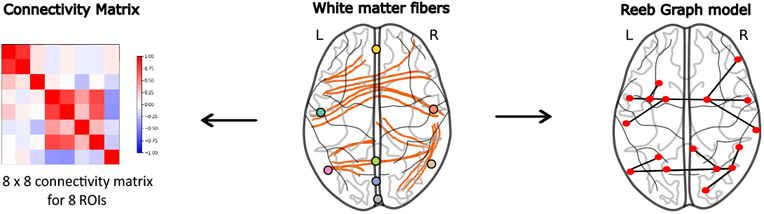
Modeling of white matter fibers using DTI. Connectivity matrix represents interconnection between different regions of interest (ROIs). Reeb graph allows the relevant geometric and topological structure of the streamlines to be recovered that is ignored in the connectivity matrix.

A key limitation of both connectome and tract-specific analysis methods is that the underlying topology of white matter bundles as they traverse the brain is lost in the dimensionality reduction process. Moreover, alterations in bundles due to neurological disorders cannot be localized since streamlines are usually lumped to a single cluster or a given region of the brain. Hence, to computationally model effects such as the increase in sparsity of bundle specific white matter fibers with the progression of Alzheimer’s disease (AD) [20] or the disruptions caused due to stroke [7], a hypothesis-driven computational geometry approach is necessary. Towards that end, a Reeb graph-based approach in [21,22] discovers the branch and merge structure of the streamlines for a topological understanding of white matter fibers. However, basic Reeb graph computation fails to account for false positives and is not robust to the noise that the tractography methods introduce. Noise in this context refers to streamlines that are geometrically implausible due to curvature overshoot; short length because of premature termination; and spatially distant due to connection density biases [31]. We address the challenges discussed above in this paper. Our main contributions are summarized as follows:

1. We develop a robust Reeb graph model that mitigates the effects of noise and false positives in streamlines,
2. We validate the robustness of our approach using ISMRM datasets, and
3. We quantify the white matter degeneration and increase in sparsity in progressive AD using the ADNI dataset.

## 2 Method

### 2.1 Problem Setup

White matter fibers in brain space are represented as a set of spatial trajectories in 3D Euclidean space such that each streamline is a trajectory *T* = {*p*_1_, *p*_2_, …, *p*_*m*_} where *p*_*i*_ ∈ ℝ^3^ are an ordered sequence of points and *m* is the total number of points. Similarly, a subtrajectory of a parent trajectory *T* starting at *s* and ending at *t* is represented by a consecutive subsequence of points {*p*_*s*_, *p*_*s*+1_, *p*_*s*+2_, …, *p*_*t*_|*p*_*i*_ ∈*T*}. A common behavior of streamlines is that they start from one brain region, merge with other streamlines into bundles, and then split towards the end into other brain regions. Intuitively, we can assume that if a continuous portion of a set of fibers are close together then they share a common behavior. In this paper, we model the topology of these streamlines by formulating a problem of computing a robust Reeb graph model ℛ (*V, E*) given a set of trajectories ℐ= {*T*_1_, *T*_2_, …, *T*_*n*_}, where *n* is the number of trajectories. The vertices *V* of the Reeb graph encode the termination, merge, or split of trajectories and the set of edges *E* are the group of subtrajectories bundled together.

### 2.2 Definition of Bundling Structure

Consider a pair of points *p*_1_, *p*_2_ ∈ ℝ^3^, we define *d*(*p*_1_, *p*_2_) = ‖*p*_1_ − *p*_2_‖_2_ as the Euclidean distance between the two points where ‖·‖_2_ is the Euclidean norm. The two points are *ϵ*-connected if *d*(*p*_1_, *p*_2_) ≤ *ϵ*. Two trajectories (or subtrajectories) *T*_*i*_ and *T*_*j*_ are *ϵ*-connected if ∀*p*_*i*_ ∈ *T*_*i*_, ∃ *p*_*j*_ ∈ *T*_*j*_ such that *p*_*i*_, *p*_*j*_ are *ϵ*-connected. Given ℐ and a distance parameter *ϵ*, bundling structure of a set of trajectories is modeled using a Reeb graph of the *ϵ*-connected trajectories [21,22]. We define:

#### Bundle

A Bundle denoted by *B* is a set of subtrajectories that consists of at most one subtrajectory from each trajectory *T*. Two bundles *B* and *B*′ are adjacent if ∃ *U* ∈ *B* that is continued as *U* ′∈ *B*′, where *U* and *U* ′ are the subtrajectories of *T*. *B* is *ϵ*-bundle if any two subtrajectories in *B* are *ϵ*-step connected i.e, connected by a sequence of *ϵ*-connected subtrajectories. An *ϵ*-bundle *B* is max-width if no other possible *ϵ*-bundle of ℐ intersects *B* and contains a super set of the trajectories represented by *B*.

#### Max-width *ϵ*-bundle partition

Given a set of max-width *ϵ*-bundles, a max-width *ϵ*-bundle partition is defined as 𝒫= {*B*_1_, *B*_2_, …, *B*_*l*_} such that every point in ℐ is assigned to a bundle. To compute 𝒫, we solve a subtrajectory clustering problem as discussed in [3,21] by maintaining a spatially dynamic graph *G* representing the *ϵ*-connected relation. The graph *G* changes with increasing spatial position: when trajectories are *ϵ*-connected we insert new edges into *G*, and when *ϵ*-disconnected we remove edges. The connected components in *G* are the bundles that are assigned to each point in ℐ giving 𝒫. Finally, the Reeb graph ℛ_0_ for 𝒫 is an undirected graph where each edge represents a max-width *ϵ*-bundle and each pair of edges *e*_*i*_ and *e*_*j*_ are connected with a vertex if *B*_*i*_ and *B*_*j*_ are adjacent bundles as shown in Fig. 2. Note that the Reeb graph is monotonous with respect to the distance parameter *ϵ* i.e. if *B* is a bundle for a given *ϵ*, then for any *ϵ*′ *> ϵ*, ∃ *B*′ ⊇ *B*. The monotonicity property ensures consistency between models at different levels of detail and is important in obtaining coarse-grained models for the streamlines.

**Fig. 2.**
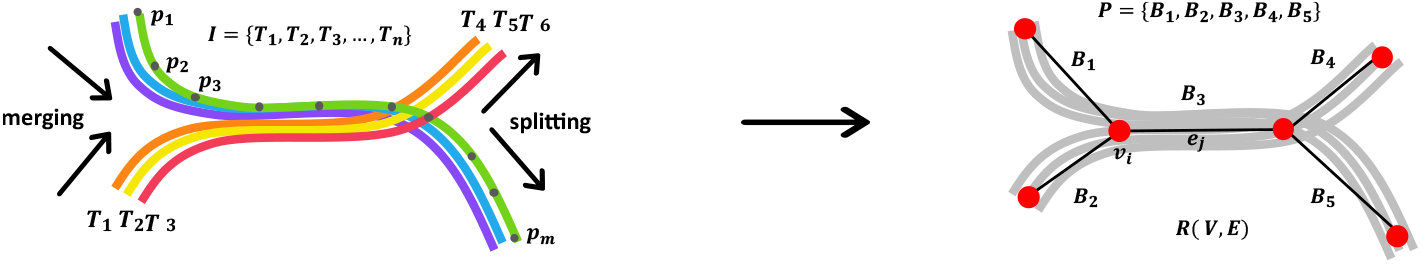
Bundling structure of streamlines. An example showing the branching structure of the streamlines and the Reeb graph where nodes encode the merge, split and termination characteristics.

### 2.3 Robust Definition of Bundling Structure

The bundling structure discussed above is susceptible to noises and errors in the streamlines as discussed in Sec. 1. It has been shown that tractography techniques suffer from a large number of false positives due to reconstruction of streamlines that do not correspond to real anatomical bundles [16]. Hence, we propose a robust Reeb graph model to detect and discard outlier streamlines based on their geometrical properties such as length, density, size, and branching. For example, minor interruptions of length *<*2 intervals in the streamlines would be insignificant when modeling the bundling structures at the 20 interval length scale. Hence, the amount of interruptions that may be considered insignificant is a latent parameter dependent on the modeling abstraction and the hypothesis underlying the model. We introduce a parameter *α* ≥ 1 to model this measure of topological persistence. Therefore, in the robust Reeb graph model, the precise events with length at most *α* are ignored. So, if a pair of trajectories are *ϵ*-connected or disconnected for less than *α* interval then the associated event is reversed. Formally, given an input image ℐ and its max-width bundle partition 𝒫, we define:

#### α-relaxed ϵ-connected bundles

Two trajectories are *α*-relaxed and *ϵ*-connected at *k* if and only if they are *ϵ* connected at some *k*′ ∈ [*k* − *α/*2, *k* + *α/*2].

Finally, to control the thickness of the bundles, we introduce a parameter *δ* to model the minimum size of the bundles in the robust Reeb graph. This allows us to model the significant bundles whose size is more than *δ* trajectories and all other bundles are ignored. To sum up, we say that a bundle *B* is robust if it is a bundle according to the definition in Sec. 2.2 with its components replaced by *α*-relaxed *ϵ*-connected components and |*B*| *> δ*. Note that the robust Reeb graph also manifests the monotonicity property in the parameters *α* and *δ*. That is, a bundle *B* in an interval *S* remains a bundle in *S* on decreasing *δ* or *α*.

##### Theorem 1.

*For a given ϵ and a set of trajectories* I, *where each trajectory consists of at most μ points, Reeb graph* ℛ_0_ *can have O*(*μn*^2^) *vertices*.

*Proof*. Consider a pair of trajectories *T* and *T* ′. *T* ′ is *ϵ*-connected to *T* during at most one spatial interval which yields two vertices in ℛ_0_. A pair of trajectories produce *O*(*μ*) vertices when connected and disconnected for at most one interval. For all possible pairs *n*^2^, this gives a total of *O*(*μn*^2^) vertices. This bound is tight in the worst case.

##### Theorem 2.

*For a given ϵ, α, and a set of trajectories ℐ, where each trajectory consists of at most μ points, the robust Reeb graph* ℛ *can have* 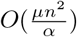 *vertices*.

*Proof*. Consider two trajectories *T* and *T* ′. *T* ′ is *ϵ*-connected to *T* during at most *α* spatial interval which yields to two vertices in ℛ. A pair of trajectories thus produces 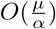 vertices. There are *n*^2^ possible pairs giving a total of 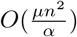 vertices. This bound is tight in the worst case.

The last step of our method is to compute the robust Reeb graph model, (*V, E*). It is an undirected graph where the set of edges, *E*, represent the set of max-width *α*-relaxed *ϵ*-connected bundles and two edges *e*_*i*_ and *e*_*j*_ are connected with a vertex, *v* ∈ *V*, if their corresponding bundles *B*_*i*_ and *B*_*j*_ are adjacent. The time complexity for computation of this model is 𝒪(*N* log *N*), same as that of the non-robust model.

## 3 Results

### 3.1 Validation

To demonstrate that the Reeb graph model is practical and indeed captures the bundling behavior of the trajectories, we implement our algorithm for both robust and non-robust cases. We use two types of data sets to evaluate our method: synthetic data (included with source code [1]) and the ISMRM dataset [15]. We illustrate the behavior of our method by providing qualitative verification of the bundles that our algorithm discovers in the presence of noise and false positives in Fig. 3. We account for the noisy streamlines by making one fundamental assumption that fibers are naturally organized as bundles in the brain. Therefore, curvature overshoot and short length streamlines are eliminated using the persistence parameter *α* while the isolated streamlines are excluded using the bundle size parameter *δ* as shown in Fig. 3A. Several complex fiber configurations are observed in tractographs [2] as false positives where streamlines cross, diverge, and turn resulting in crossing, fanning, and kissing characteristics as illustrated in Fig. 3B. Such confounding patterns can be included or excluded in our model using the robustness parameters: *α* is set to the length of the small encounters to handle the crossing and kissing patterns while *δ* is set to the size of the sparse bundle at the end to account for the fanning configuration.

**Fig. 3.**
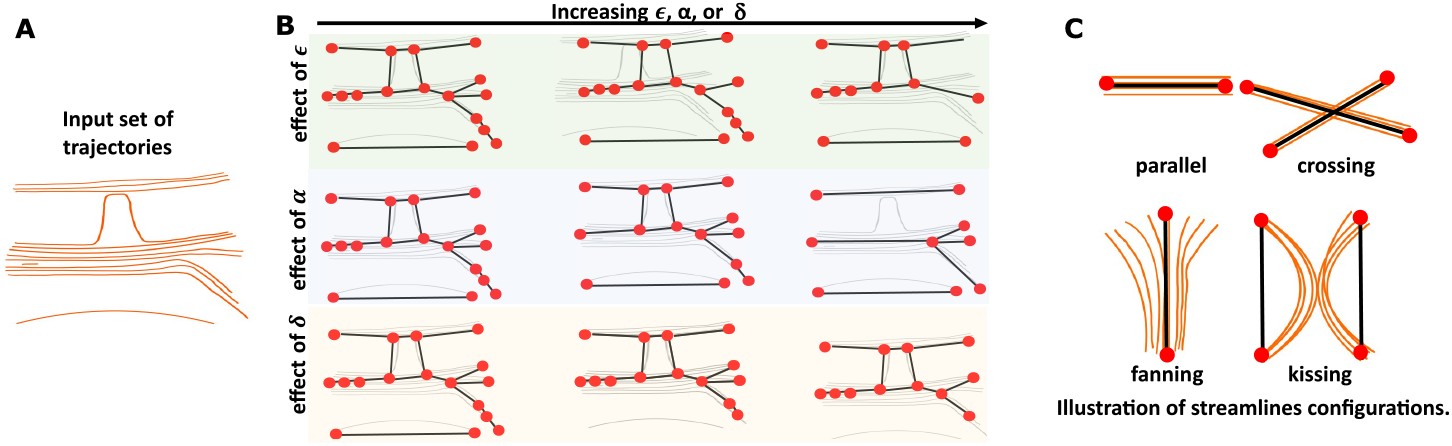
Validation of the robust Reeb graph using synthetic data. (A) Input set of streamlines generated using GeoGebra [4]. (B) Tuning *ϵ, α*, and *δ* obtains generalized views of the grouping structure and handles false positives. (C) Common examples of false positives observed in tractography and their robust models.

#### ISMRM dataset

This publicly available dataset [14] consists of 25 major tracts that are representative of the challenges in human brain imaging *in vivo*. Tracts span different shapes, lengths, and sizes that help us in validating the output of our proposed method and analysing the effect of granularity and robustness parameters (*ϵ, α*, and *δ*). Fig. 4 shows the Reeb graph representation of three well known neurological structures from the ISMRM dataset. The robust Reeb graph model detects and handles the morphology of the white matter tract configurations due to branching and local ambiguities such as crossing, kissing, and fanning. We visually inspect the maximal groups identified by our algorithm and verify the resulting model by using domain expertise. As an illustration, in all four examples of fiber tracts shown in Fig. 4, no interruption is expected at the middle segments while a branching structure is expected at the end. Our method captures these branching structures. Furthermore, our model discovers the known termination regions and assigns them nodes (and hence, locations in brain space) in the Reeb graph.

**Fig. 4.**
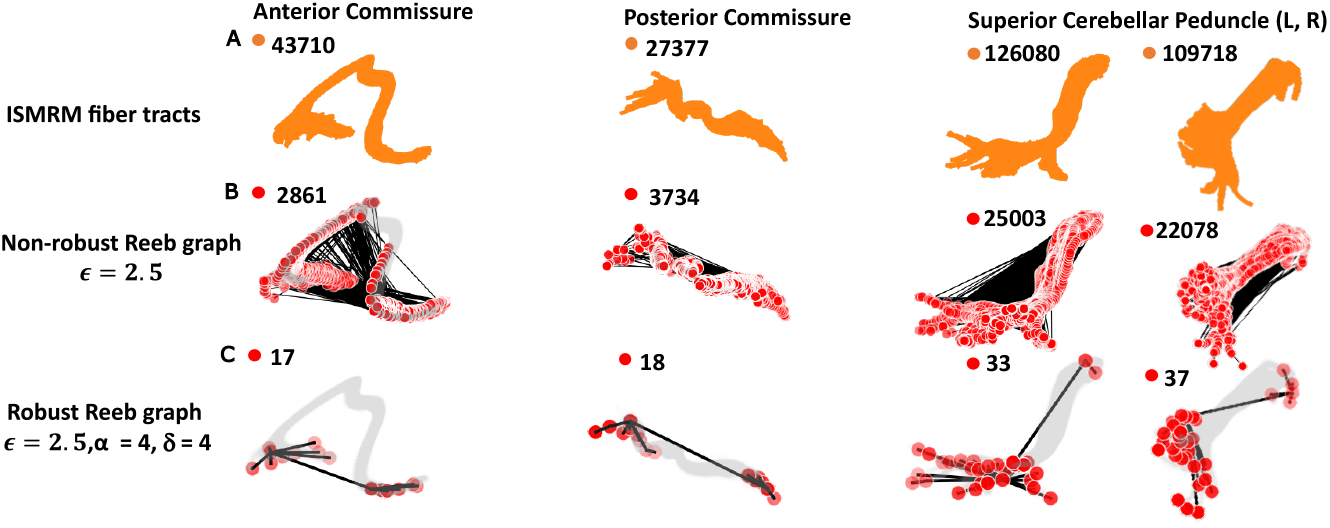
Validation using ISMRM dataset. (A) Streamlines from the dataset. (B) Non-robust Reeb graphs for the streamlines. (C) Robust Reeb graph representation. We can observe that the non-robust Reeb graph shows extraneous nodes and edges due to noise while the robust graphs identify anatomically correct structures.

### 3.2 Modeling Alzheimer’s Disease Progression

We investigate the anatomical changes in the ADNI dataset [17] comprising of AD and normal controls with scans from multiple visits.

#### Data Collection and Pre-Processing

Brain DTIs from 31 subjects with age 75 – 85 years are used in this study. All the subjects are selected under the given imaging protocols: the scans are from the same manufacturer, GE Medical Systems and the magnetic field strength is 3 Tesla. A total of 62 pre-processed scans with Eddy current correction are downloaded, two scans for each subject with ∼1 year gap between visits. The ICBM152 AAL2 (Automated Anatomical Labeling, AAL [23]) brain atlas is used to divide the brain of each subject into 120 brain regions in total. 10 brain regions relevant to AD and 8 regions known to be preserved in AD are used as ROIs [25,18] as shown in Fig. 5. For each ROI, the diffusion data is reconstructed in the Montreal MNI (Montreal Neurological Institute) space using q-space diffeomorphic reconstruction [27]. A deterministic fiber tracking algorithm [28] is used to compute fiber tracts.

**Fig. 5.**
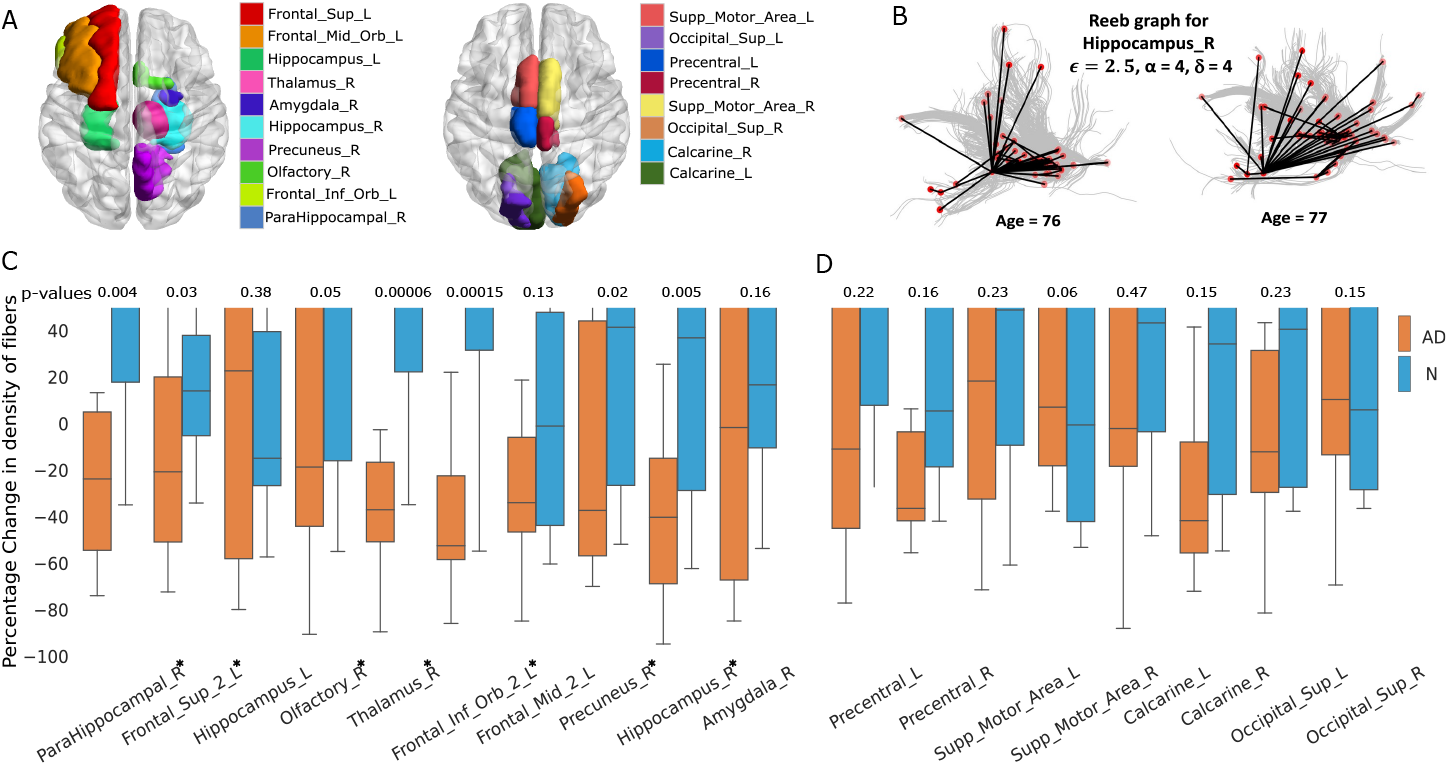
Reeb graph analysis of brain ROIs. (A) Relevant and preserved brain regions with respect to AD. The brain regions are visualized using BrainNet Viewer [26]. (B) Robust Reeb graph models with age for HippocampusR. Highlighted nodes in the Reeb graph (right) show sparse and small length bundles. (C, D) The box plot of *Δρ*^*^ for AD and normal control group for relevant and preserved brain regions respectively. Relevant brain regions (C) show severe decrements in the fiber density (ROIs marked with *) while we observe mild changes in the preserved brain regions (D).

#### Weighted Reeb Graph

Bundles in a given image are associated with edges in the Reeb graph and can be labeled with domain-specific information. Here, we define fiber density as an anatomical feature for Alzheimer’s disease progression.

##### Fiber density (*ρ*)

For a given input image ℐ and its max-width bundle partition 𝒫={*B*_1_, *B*_2_, …, *B*_*k*_}, we define fiber density for each bundle *B* denoted by *ρ* as *ρ* = |*B*| */n*, where |*B*| is the total number of trajectories in bundle *B*.

To quantify the sparsity of the model, we use *ρ* as the edge weights. Further, we compute percentage change in fiber density as *Δρ* = (*ρ*_*a*_ − *ρ*_*b*_)*/ρ*_*a*_ × 100, where *ρ*_*a*_ and *ρ*_*b*_ are fiber density at age *a* and *b* respectively and *a < b. Δρ*^*^ is the percentage change in the fiber density for the maximal width bundle *B*_*m*_ such that *m* = arg max_*i*_ |*B*_*i*_|. We use *Δρ*^*^ to quantify the fiber disruptions and localize the maximal robust bundle. We hypothesize that the control group will show mild white matter disruptions while AD subjects will show a pattern of disrupted connections. The hypothesis is tested using the box-plot shown in Fig. 5C,D. The results suggest that the fiber density decrements in AD patients with age is greater than that in control subjects. Moreover, the differences are statistically significant (*p <* 0.05) for 7 out of 10 relevant brain regions while insignificant for all the brain regions known to be unaffected by AD. Possible reasons for relevant regions (Hippocampus L) not showing significant p-value could be the issues of data quality or choice of parameters. Additionally, decrements in length of the bundles may also be used as a feature for AD progression modeling since it represents damaged fiber tracts. By mapping the Reeb graph nodes to the 3D brain space, our method localizes the signature of white matter disruptions in different tracts and their associated cortical regions.

## 4 Conclusion

We present a robust Reeb graph-based modeling of white matter fibers. We demonstrate that the proposed approach captures the fiber branching behavior in the presence of noise and false positives, and is further validated on synthetic and real data. Results on the ADNI data show that the method can quantify the fiber disruption in Alzheimer’s population with aging.

The choice of parameters in our method is dependent on the hypothesis for which the Reeb graph is built. For example, a detailed view (dense graph) can be obtained to compare and contrast tractography methods that generate the streamlines. Future work in this direction includes quantifying robustness in tractography. On the other hand, for disease study, a coarse model (sparse graph) can be computed such that the false positives are suppressed to quantify the disease-relevant features. Another interesting research area is to link the structural model of streamlines to the behavioral model of fMRI to gain new insights into structural and functional connectivity relationships.

